# Olfactory combinatorial coding supports risk-reward decision making in *C. elegans*

**DOI:** 10.1101/2024.06.19.599745

**Authors:** Md Zubayer Hossain Saad, William G. Ryan V, Chelyan A. Edwards, Benjamin N. Szymanski, Lana Awa, Jenna Kaake, Alexander Martin, Aryan R. Marri, Lilian G. Jerow, Robert McCullumsmith, Bruce A. Bamber

## Abstract

Olfactory-driven behaviors are essential for animal survival, but mechanisms for decoding olfactory inputs remain poorly understood. We have used whole-network Ca^++^ imaging to study olfactory coding in *Caenorhabditis elegans.* We show that the odorant 1-octanol is encoded combinatorially in the periphery as both an attractant and a repellant. These inputs are integrated centrally, and their relative strengths determine the sensitivity and valence of the behavioral response through modulation of locomotory reversals and speed. The balance of these pathways also dictates the activity of the locomotory command interneurons, which control locomotory reversals. This balance serves as a regulatory node for response modulation, allowing *C. elegans* to weigh opportunities and hazards in its environment when formulating behavioral responses. Thus, an odorant can be encoded simultaneously as inputs of opposite valence, focusing attention on the integration of these inputs in determining perception, response, and plasticity.

## Introduction

Animals must detect and respond to chemical signals in the environment to find food, avoid hazards, and find mates. Olfactory systems balance two competing priorities to optimally represent the external chemical space. First, they must respond rapidly and robustly to specific compounds which elicit innate responses important for survival. These include predator or parasite odors (eliciting avoidance), and pheromones (eliciting social and/or reproductive responses) (*1–4*). Such stimuli are sensed using narrowly tuned odorant receptors in the periphery, hard-wired to dedicated circuits for specific behaviors (labeled line coding). Second, olfactory systems must be versatile, capable of detecting and discriminating millions of different environmental odorants. This versatility is achieved using broadly-tuned odorant receptors, which activate in different combinations depending on the chemical structure of the odorant, enabling millions of different odorants to be discriminated using, at most, a few thousand receptors (*5*). These combinations are decoded centrally to identify the odor (combinatorial coding)(*4*). Combinatorial coding has been regarded as the predominant mechanism for olfactory coding (*6, 7*), but examples of labeled line-like olfactory coding are increasingly being found (*8–12*), suggesting that olfactory coding is a hybrid of labeled line and combinatorial mechanisms (*1*). Labeled line coding is simpler, and probably evolved first, with animals responding to specific environmental cues using specific receptors. Combinatorial coding likely emerged later as the narrowly tuned receptor genes underwent duplication and divergence to diversify ligand binding properties. Presumably, the central decoding circuitry then evolved to take advantage of these new sources of environmental information (*1, 13*). The initial steps in olfactory coding are relatively well studied, with each olfactory receptor neuron expressing 1-2 olfactory receptors (*14, 15*). Inputs from olfactory receptor neurons expressing the same receptor converge on a single glomerulus in the olfactory bulb (mammals) or antennal lobe (insects), and the glomeruli project to higher processing centers in the brain (*16*). Recent studies are beginning to unravel the neural circuitry by which olfactory inputs generate perception and responses (*17–20*). However, we still lack a clear understanding of how combinatorial inputs are decoded, and how combinatorial pathways and labeled line pathways may functionally interact.

We have turned to *C. elegans* to study the downstream processing of olfactory inputs. The small size of the *C. elegans* nervous system and the availability of techniques for pan-neuronal activity recording now make it possible to study the neural basis for sensory-driven behaviors at whole-network scale (*21–31*). Olfaction in *C. elegans* has classically been viewed as a labeled-line coding system rather than a combinatorial system, where specific subsets of chemosensory neurons respond to attractive odorants and drive positive chemotaxis (AWC, AWA), or respond to repulsive odorants and drive negative chemotaxis (ASH, AWB, ADL) (*32–38*). However, it was recently shown that individual odorants activate multiple classes of olfactory neurons, and individual olfactory neurons respond to a wide range of odorants (*39*). This responsiveness profile is the hallmark of combinatorial coding, even though there are many fewer chemosensory neurons in *C. elegans*, and these neurons must express a much wider repertoire of olfactory receptors than their mammalian counterparts. The striking parallels in olfactory coding between animals as divergent as nematodes, insects, and mammals suggests that evolution may have converged on a few optimal solutions for sensing and responding to external odor landscapes (*40*).

We have performed an in-depth analysis of a single odorant, 1-octanol (1-oct), which is encoded combinatorially (*39*), and is well-known to stimulate aversive behavior (*32, 33, 38, 41*). We show that 1-oct stimulation concurrently activates separate attractive and repulsive sensory afferent pathways in the nervous system. These pathways converge to control at least four aspects of locomotion at the behavioral and circuit levels, with the balance between attraction and repulsion emerging as the critical factor determining the overall behavioral output. Furthermore, this coding strategy provides a simple framework for modulating sensory-driven responses to produce context-appropriate behavior.

## Results

### 1-octanol repulsion in microfluidic arenas

1-oct avoidance in *C. elegans* has been intensively studied because it is strongly modulated by nutrition state and monoamine/neuropeptide signaling cascades (*34, 42–46*). Our goal was to investigate the network basis of 1-oct repulsion using the NeuroPAL system (*30*), which allows simultaneous whole-brain Ca^++^ imaging with unambiguous neuronal identification, coupled with quantitative analysis of 1-oct aversive behavior. 1-oct has previously been utilized as a volatile stimulus (*33, 34*), or in a gradient on agar plates (*47, 48*), while NeuroPAL recordings are performed in aqueous solution in microfluidics chips under confocal microscopy (*30*). To better match the conditions for NeuroPAL recordings and behavioral assays, we first investigated 1-oct repulsion in microfluidics arenas (*49–51*), which gave us precise control over the concentration and spatiotemporal characteristics of the 1-oct stimulus. Using a simple two-chamber arena (Fig. 1A), we demonstrated that worms were repelled by a saturating concentration of 1-oct in S-basal (2.2 mM, corresponding to a 3.75 x 10^-4^ dilution; Fig. 1B-D). In our initial trials, we added 0.01% w/v xylene cyanol (0.1 mg/mL) to the 1-oct solution to visualize the interface between the stimulus and control buffer. However, as another study reported (*51*), worms were significantly attracted to xylene cyanol, introducing an unwanted stimulus into the assays. We therefore optimized methods to maintain a stable interface between odorant and buffer chambers without the need for dye (see Methods). We assayed worms between 20 and 40 minutes after removal from food, during area-restricted search behavior (*52*), and calculated average chemotaxis indices (CIs) between 30 and 40 minutes (see Methods, and Fig. 1D, E). The -0.7 CI observed in the trial shown represents strong repulsion. The average CI over n=5 was -0.27±0.24 (SEM).

**Fig. 1.**
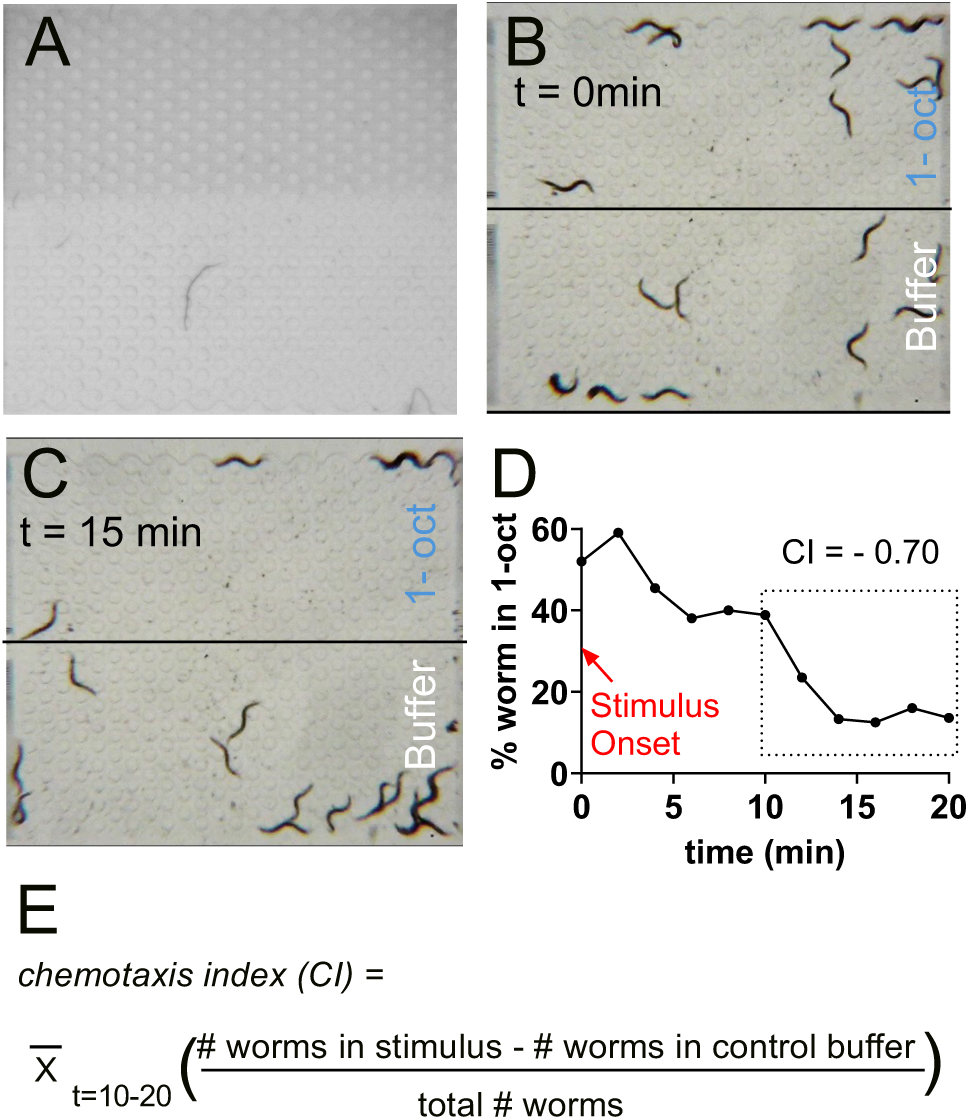
1-oct chemotaxis in microfluidic arenas. (**A**) Microfluidics arenas were configured with two zones of equal area, one containing a 1-oct solution and the other containing buffer alone (1-oct zone contains 0.01% xylene cyanol dye for visualization here; assays are run without dye). (**B, C**) Typical distribution of worms when first exposed to 1-oct (B), and after 15 min of 2.2 mM 1-oct exposure (C). (**D**) Time-course of 1-oct chemotaxis for the trial shown in B, C (**E**) Calculation of chemotaxis index, averaging worm counts every 2 minutes from 10-20 minutes after introduction of 1-oct solution.

### Network-wide response to repulsive olfactory stimulation

To investigate the neuronal basis for the observed 1-oct aversive behavior, we performed network-wide Ca^++^ imaging during 1-oct stimulation and identified the responding cells by imaging the neuronal landmarks encoded by the NeuroPAL fluorescent reporters (Fig. 2A). Worms were immobilized in ‘olfactory’ microfluidics chips (*53*) for microscopy. Previous whole-brain studies have shown that certain neurons undergo spontaneous transitions between low and high activity states correlating with spontaneous reversals during foraging, notably the reverse command neurons AVA, AVE, AVD, AVB, and the RIM neuron (*23, 27, 54–59*). We therefore used a recording/stimulus protocol consisting of a 6 min pre-stimulus period, followed by 6 30s applications of 2.2 mM 1-oct, spaced 30s apart, to help distinguish between spontaneous transitions and potential sensory-evoked transitions within this neuronal population. We observed dynamic activity patterns in dozens of neurons before and during 1-oct stimulus (Fig. 2B-D). Consistent with previous studies, AVA, along with AVE, AVD, RIM, AIB, and AIZ showed spontaneous state transitions, with the AVB forward command interneuron and the AIY interneuron showing mirror image patterns (i.e. AVB/AIY inactivated when AVA activated and vice versa). Typically, 3-4 transitions were observed in a 12 minute recording. These transitions took place during both the pre-stimulus and stimulus periods (Fig. 2D, 3A) and showed no obvious relationship with 1-oct application or withdrawal. These transitions are related to network state changes which drive spontaneous reversals during foraging in freely moving worms (*60*). Immobilization and anesthetization, necessary for confocal imaging, distort certain aspects of these motor command sequences compared to freely moving worms executing the motor commands and receiving proprioceptive feedback. However, the intrinsic motor programs remain intact under these conditions (*60*).

**Fig. 2.**
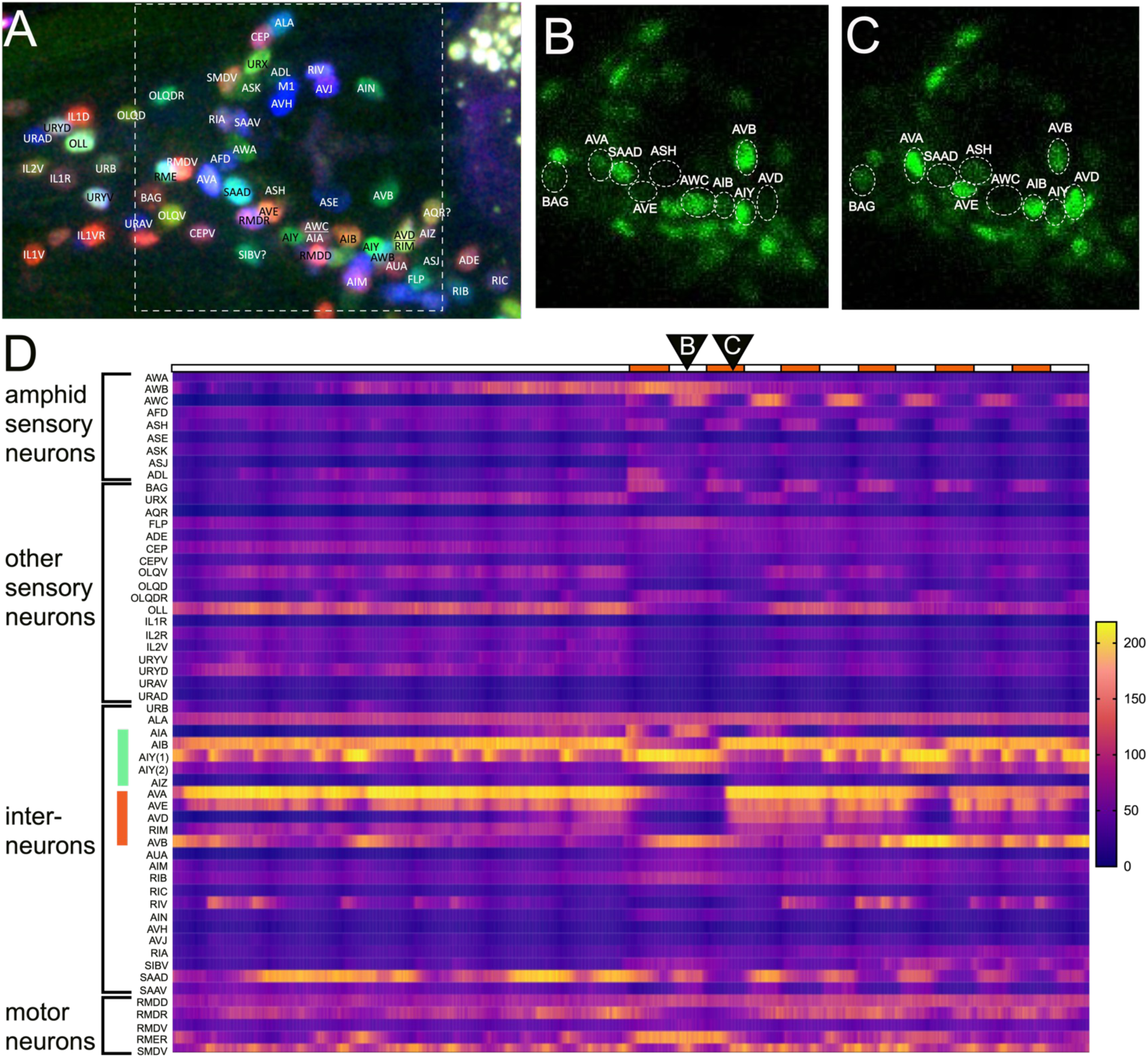
Network recording of 1-oct (2.2 mM) response. **(A)** Representative image of a NeuroPAL worm immobilized in a microfluidics device visualized using the mTagBFP2, CyOFP, and mNeptune2.5 reporters for neuronal identification (*22*). (**B, C**) Pan-neuronal nGCaMP6s signal for the region indicated by the box in (A) during 1-oct stimulation, showing 1-oct offset (B) and onset (C). Specific time points in the recording for (B) and (C) are indicated by the black triangles in (D). Selected sensory neurons and interneurons are indicated by dashed white circles. (**D**) Heat map of activity profiles for 56 identifiable neurons. Raw (i.e. non-normalized) fluorescence values shown. Stimulation time-course shown on top row (white is buffer, orange is 1-oct). Green and orange vertical bars at left indicate first-layer and locomotory command neurons (+RIM), respectively.

Sensory signals were also prominent, in both amphid and non-amphid chemosensory neurons. ASH and BAG showed strong consistent responses correlating with stimulus onset (‘ON’ responses). AWB also showed an ON response but also had high levels of spontaneous activity. ADL showed a strong ON response to the initial stimulus but was strongly desensitized in subsequent applications (Fig. 2D and 3B). ASH, AWB, and ADL were previously identified as the main neurons mediating 1-oct aversive responses (*33, 34*). 1-oct responses were also observed in AWA, ASE, ASK, ASJ, and URX (Fig. 2D, 3B), consistent with the observation of Lin et al, showing that 1-oct activates sensory neurons promiscuously (*39*). Significantly, we observed a consistent strong 1-oct response in AWC neurons (both AWC^OFF^ and AWC^ON^; Fig 2D, 3B). AWC activity decreased upon 1-oct addition and rebounded upon 1-oct withdrawal, in a typical ‘OFF’ neuron pattern (*61*), opposite to ASH (Fig. 2B-D, 3C). This was surprising, first, because AWC is usually considered an attractive neuron, not expected to respond to a repulsive odorant, and second, because the observed AWC pattern predicts that AWC should drive attraction to 1-oct (*38, 47, 61, 62*). 1-oct has thus far only been described as a repellent for *C. elegans*, with no attractive properties (*32, 34, 38, 48*). Strong sensory-driven activity was also observed in AIA and SAA interneurons, suggesting these two neurons may receive substantial inputs from the sensory neurons (Fig. 2D, 3C). Apart from AIA, the other ‘first layer’ interneurons AIB, AIY, and AIZ did not show detectable sensory-driven activity (Fig. 3A), despite their extensive anatomical connections with sensory neurons, suggesting these interneurons may be receiving their principal functional inputs from elsewhere in the circuit.

**Fig. 3.**
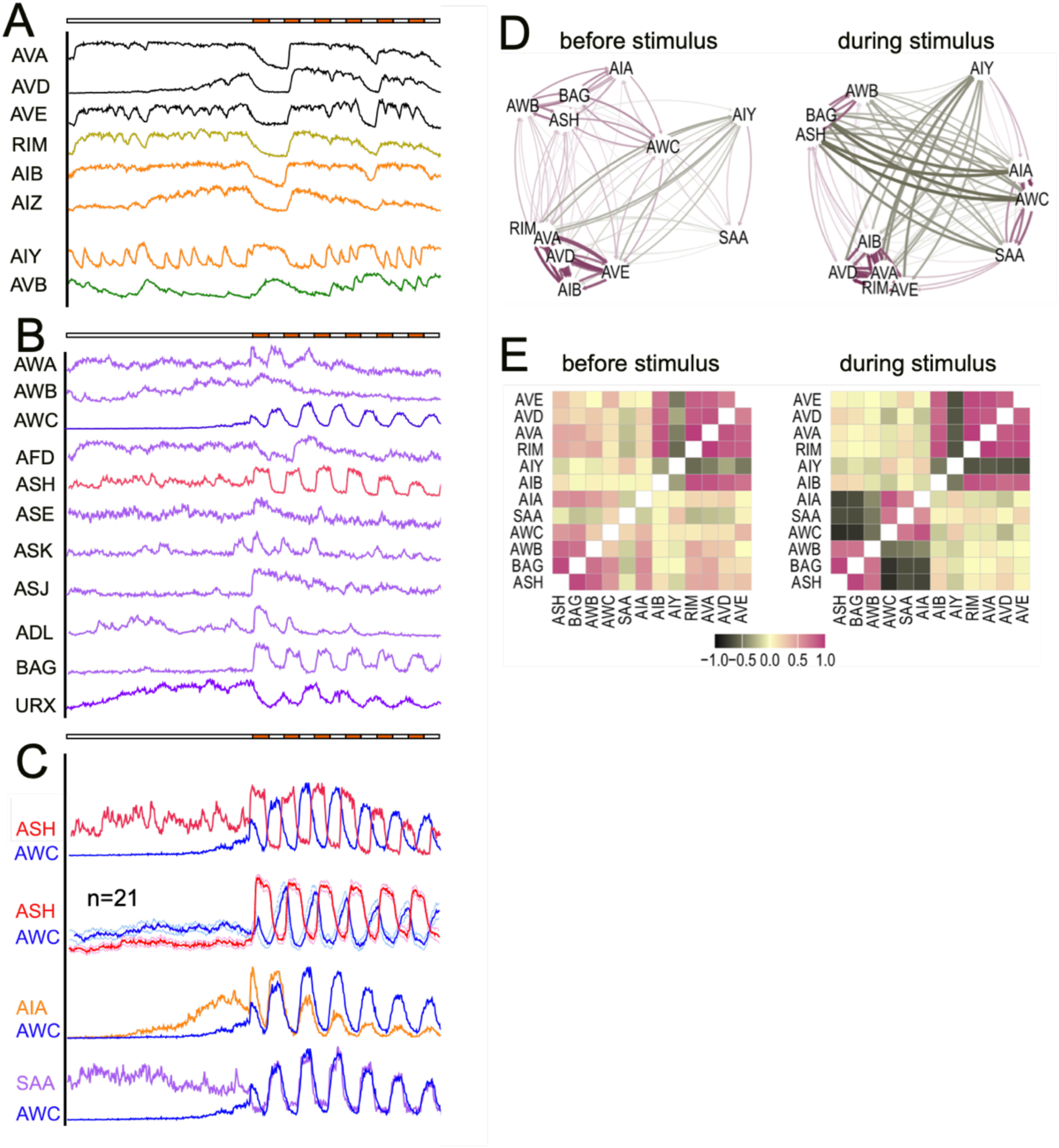
Neuronal activity correlations. **(A-C**) Neuronal activity patterns from worm shown in Fig 2. (**A**) Stochastic activity state transitions of locomotory command and associated interneurons, with AIY and AVB showing opposite activity patterns to AVA, AVD, AVE, RIM, AIB, and AIZ. (**B**) Sensory neuron activity patterns. (**C**) Selected neuron pairs highlighting the anti-correlated activity patterns of AWC and ASH (top two traces; averaged AWC/ASH data from 21 worms shown to highlight consistency of anti-correlation between these two neurons), and the correlated activity patterns of AWC with AIA (middle) and SAA (bottom). (**D, E**) Activity correlations for 12 selected neurons based on data from 5 worms. (**D**) Graph representation of activity correlations before stimulus (minutes 0-6, left) and during stimulus (minutes 6-12, right). Neurons pairs with the strongest positive correlations are closest to one another and connected by the darkest purple lines; neurons with the strongest negative correlations are furthest apart and connected by the darkest green lines. (**E**) Relationships from (D) presented as a correlation matrix using the same color code.

To better understand how sensory inputs may pattern global network activity patterns, we performed correlation analysis on selected data sets in which we were able to identify 12 key neurons with well characterized roles in sensory motor coupling (*38, 52, 61*). To detect the sensory signature, we compared neuronal correlation patterns in the first 6 minute interval (prior to stimulation), and the second 6 minute interval (where stimulus was applied and removed for alternating 30s periods) (Fig. 3D, E). We observed the expected increases and decreases in correlation among the sensory neurons (i.e. ASH and BAG became correlated, and anti-correlated with AWC). We did not observe any increased correlation between the motor command interneurons and sensory neurons, consistent with a lack of observable sensory-driven activity in the motor command interneurons. However, the degree of activity cross-correlation among AVA, AVE, AVD, AIB, and RIM increased during the stimulus period, and AIB also showed increased correlation with ASH. These observations suggest that sensory information was penetrating to the locomotory command circuit level even if there was no obvious sensory entrainment of the principal neurons in those circuits (Fig. 3D, E).

Finally, we observed several neurons with activity patterns which did not correspond to the locomotory command group or the sensory-dependent group, which included SMDV, RIA, and RMDD (Fig. 2D), which are generally associated with turning behavior (*63–65*).

### The balance of simultaneous repulsion and attraction shapes the chemotactic response

The observation that 1-oct application and withdrawal cycles activate ASH and AWC in alternation predicts that these two neurons generate antagonistic inputs into the locomotory circuitry. Specifically, ASH activity varies in phase with 1-oct application, predicted to stimulate in-phase reversal consistent with a repulsive chemotactic response. AWC activity varies out of phase with 1-oct application, predicted to inhibit in-phase reversals and stimulate out of phase reversals, consistent with an attractive chemotactic response. This puzzling activity pattern led us to hypothesize that 1-oct is encoded combinatorially, simultaneously as a repellant and an attractant through two distinct and opposing afferent pathways, with the balance of these pathways determining downstream events and eventual behavioral outcome.

To test this idea, we conducted several experiments to selectively inhibit the putative repulsive and attractive afferent pathways singly, with the goal of unmasking the effects of the opposite pathway. First, to selectively diminish the repulsive pathway, we reduced the 1-oct concentration. Lin et. al. reported 1-oct concentration-response relationships for the amphid sensory neurons including ASH and other neurons implicated in 1-oct avoidance (ADL, AWB), and AWC (*39*). At sub-saturating concentrations, the responses of ASH, and to a lesser extend AWB and ADL, showed clear concentration dependence, while the ‘ON’ response of AWC did not. We extended these observations to the concentration ranges we used in our behavior and imaging experiments, measuring ASH responses, and both the AWC ‘ON’ and ‘OFF’ responses, during serial stimulation of individual worms (0.22 mM, 0.66 mM, and 2.2 mM 1-oct). ASH responses showed significant concentration-dependent increases, whereas AWC ‘ON’ and ‘OFF’ responses did not (Fig. 4A-C). In parallel, we found that lower concentrations of 1-oct (0.22 and 0.66 mM) were strongly attractive (Fig. 4D), while intermediate concentrations were largely without effect (Fig. 4D). Thus, the 1-oct chemotactic response transitions from attraction, through neutral, to avoidance, as the activity of ASH (and possibly other neurons) increases relative to AWC, highlighting the potential importance of the balance of repulsive and attractive input pathways (note that 1-oct repulsion at 2.2 mM in wild type was modest and highly variable; the solubility limit of 1-oct in aqueous solution prevented testing higher concentrations). Next, we took a genetic approach using *tax-4* and *osm-9* mutations to selectively inhibit different sets of sensory neurons. TAX-4 is a cGMP-gated ion channel required for signaling in AWC and other neurons (including AWB, BAG, ASE); OSM-9 is a TRP channel which contributes to signaling in ASH and other neurons (including ADL, AWA) (*38, 44, 66*). Loss of TAX-4 caused 1-oct to become repulsive at all concentrations (Fig. 4E), consistent with an obligatory role for TAX-4 in the putative attractive pathway. Surprisingly, loss of OSM-9 had no significant effect on CI values at any concentration (Fig. 4E), suggesting that these mutants may retain some residual ASH signaling (*38, 67*) or 1-oct repulsion may be due to multiple sensory neurons acting redundantly, some of which do not signal through OSM-9. Finally, we directly tested whether ASH and AWC, specifically, play antagonistic roles in 1-oct chemotaxis. We used chemogenetic inactivation, based on transgenic expression of *Drosophila* HisCl1 in ASH (using the *sra-6* promoter) and AWC (using the *ceh-36* promoter) (*68, 69*). Incubating these strains on histamine (His) results in acute, highly specific neuronal inactivation. Using 2.2 mM 1-oct as stimulus, His-minus controls were not significantly different from wild type for either the ASH-HisCl or the AWC-HisCl strain (despite the appearance of slight attraction under these specific experimental conditions). Importantly, the addition of His significantly skewed chemotaxis indices in the expected direction: CI values shifted positively for ASH HisCl1 worms (Fig. 4F) and negatively for AWC HisCl1 worms (Fig. 4G), consistent with reduced repulsive drive and reduced attractive drive, respectively. These findings strongly support the hypothesized antagonism between ASH and AWC in 1-oct chemotaxis. Overall, these three approaches provided evidence that 1-oct chemotaxis is driven by two distinct afferent input pathways with opposite valence in balance with one another, which can be distinguished by their differential concentration dependence, their differential usage of signal transduction pathways, and different primary sensory neurons providing the initial input (i.e. ASH for the repulsive pathway and AWC for the attractive pathway).

**Fig. 4.**
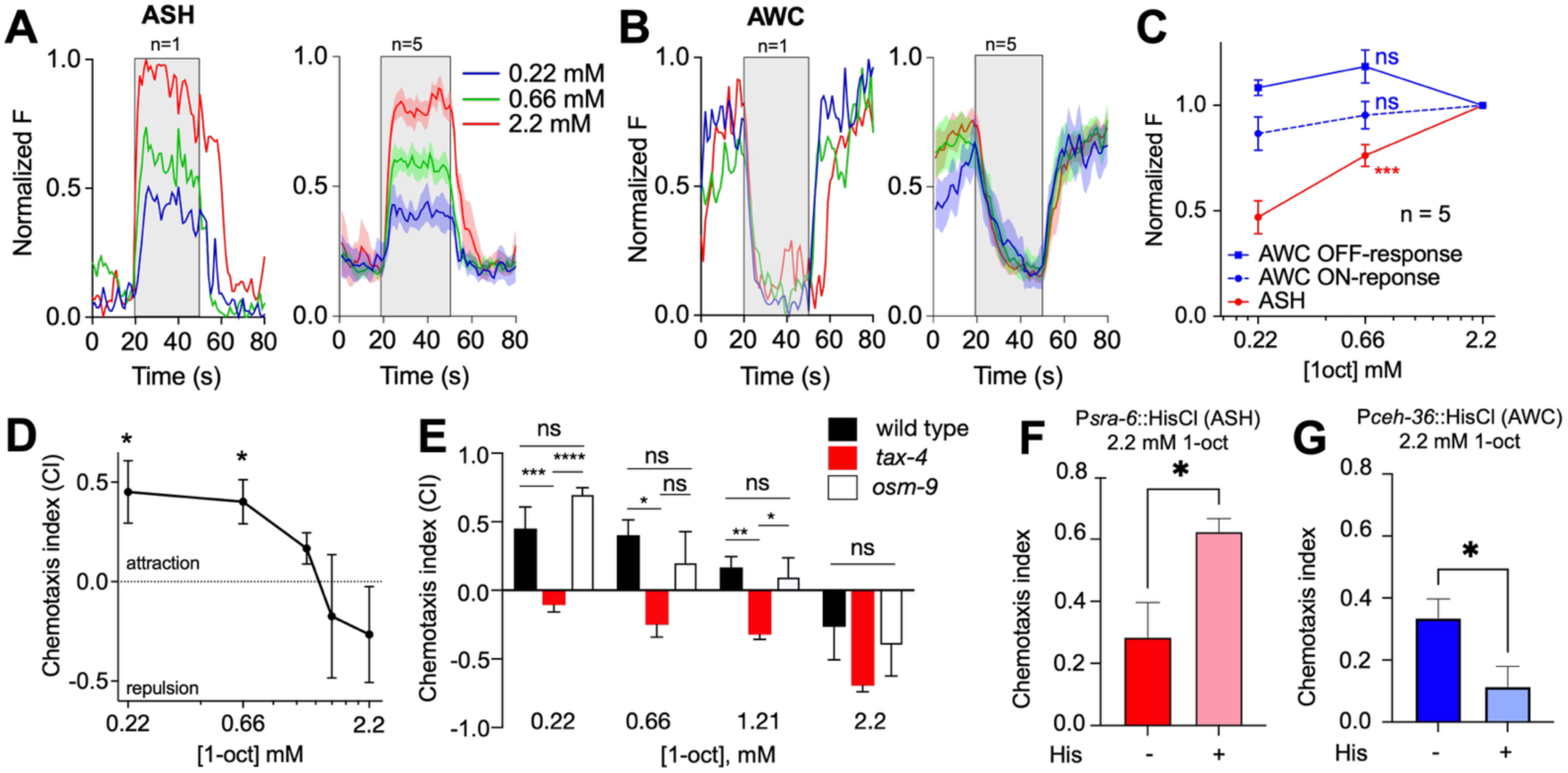
Chemotactic outcome is influenced by the balance of attractive and repulsive sensory inputs. (**A-C**) ASH ON responses (A, C) are concentration dependent, while AWC ON and OFF responses (B, C) are not, over a 10-fold concentration range from 0.22 to 2.2 mM. Representative single traces (A, B, left panels), averaged traces (A, B, right panels), and concentration-response curves (C) shown; *** P<0.001, ns P>0.05, paired ANOVA. (**D**) 1-oct chemotaxis transitions from positive (attractive) to negative (repulsive) with increasing concentration; intermediate concentrations (1.21 and 1.54 mM) result in little chemotaxis; *P<0.05 compared to 2.2 mM, ANOVA. **(E)** *tax-4* mutants are repelled by 1-oct at all tested concentrations (*P<0.05; **P<0.01; ***P<0.005; ****P<0.0005; ns P>0.05, ANOVA, n=5 for each data point). *osm-9* mutants do not differ from wild type. **(F, G)** chemogenetic inactivation of ASH, AWC: ASH inactivation increases the chemotaxis index, suggesting reduced repulsive drive (F); AWC inactivation decreases the chemotaxis index, suggesting reduced attractive drive (G) *P<0.05 *t-test*.

To understand the interaction of these co-activated afferent pathways at the level of locomotory decision making, we closely examined the behavior of worms as they encountered the interface between 1-oct solution and control buffer in microfluidic devices. We performed this analysis using the microfluidics ‘stripe’ chip (*50, 51*), which contains a central zone of 2.2 mM 1-oct flanked by zones containing control buffer. We used automated tracking (*70*) to quantify behavior upon encountering zone boundaries (i.e. reversals vs successful transits; Fig. 5A, B). For worms entering the 1-oct zone, entry transits outnumbered entry reversals by nearly 2:1 (Fig. 5B, C), which was surprising given that the overall behavioral outcome is generally avoidance under these conditions. *osm-9* mutation further depressed entry reversals, indicating that *osm-9* dependent signaling contributes to reversal in response to 1-oct in the microfluidic environment. Interestingly, *tax-4* loss-of-function strongly stimulated entry reversals, indicating that *tax-4* signaling normally suppresses these reversals, consistent with a role promoting 1-oct attraction (Fig. 5B, C). This attraction-promoting activity was also observed for worms exiting the 1-oct zone (Fig. 5B, C). Here, exit reversals strongly outnumber exit transits but in *tax-4*, the pattern is reversed, with transits outnumbering reversals by roughly 4:1. *osm-9* loss of function had no effect on exit reversals (Fig. 5C). These observations confirm that TAX-4 mediated attractive signaling significantly shapes the behavioral response to 1-oct, even though 1-oct is traditionally viewed as a strong repellent. Most importantly, they identify a locomotory transition important for chemotaxis (i.e. entry reversal frequency) in which two antagonistic sensory afferent pathways converge at a single point in time and space (Fig. 5B).

**Fig. 5.**
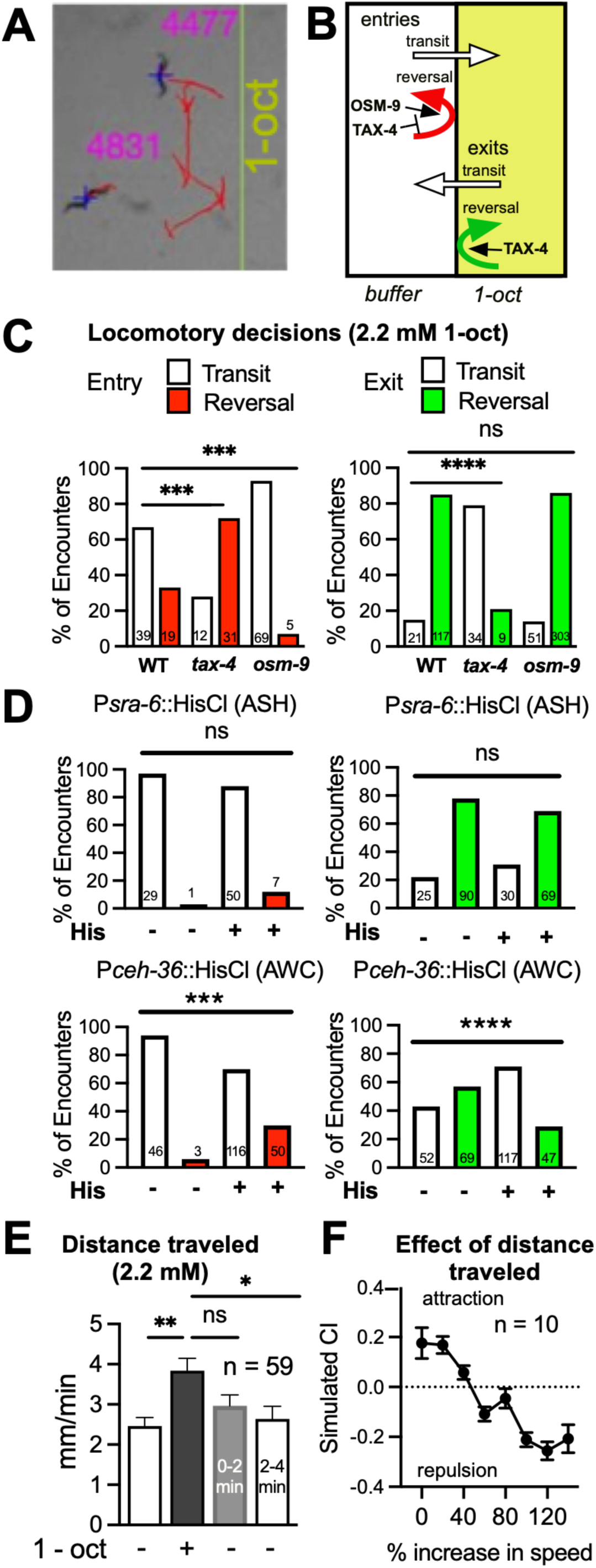
Locomotory reversals and speed modulation stimulated by 1-oct. **(A**) Example of a single worm track traced by the automated tracking software. (**B**) Worms execute reversals probabilistically at the 1-oct-buffer interface dependent on TAX-4 and OSM-9 signaling cascades. (**C**) Reversal probabilities at 1-oct-buffer interface for wild-type, *tax-4*, and *osm-9* (see panel B for summary and explanation of terms). Left panel: movement from buffer zone to 1-oct zone (‘entries’). Right panel: movement from 1-oct zone to buffer zone (‘exits’). **(D)** Reversal analysis of trials shown in 4F, G. Bars color coded as in (C). (**E**) Modulation of locomotory activity. Distance traveled in 2 minutes prior to, during, and after flooding the entire chip with 1-oct solution. (**F**) Simulation of chemotaxis outcome based on reversal probabilities (from C) and locomotory activity modulation (from E). Points shown are averages of 10 simulation trials with SEM error bars. ***, **** in C, D indicate P<0.001, 0.0001 Fisher’s Exact Test. *, ** in E indicate P<0.05, 0.01, ANOVA (Tukey’s multiple comparison test).

These findings were largely recapitulated by the chemogenetic inactivation of ASH and AWC. We analyzed reversal behavior at the 1-oct/buffer interface in the HisCl1 experiments shown in Fig. 4E and 4F (performed in the smaller two-chamber microfluidics arenas) (*71*). Inactivation of ASH had no significant effect on locomotory decision-making. However, these datasets contained very few entry reversals to begin with, so we were unable to assess the role of ASH in these events. Exit reversals were also unaffected, similar to *osm-9* trials. Importantly however, inactivation of AWC produced a very similar result to *tax-4* mutation. Entry reversals were significantly increased and exit reversals were strongly suppressed, pointing to a key role for AWC specifically in the attractive sensory afferent pathway for 1-oct chemotaxis.

These behavioral data raised a new important question: how can wild-type worms avoid the 1-oct containing zone when their reversal decisions favor entry and retention? We had noticed that contact with 2.2 mM 1-oct caused worms to rapidly and reversibly increase their locomotory activity, suggesting an additional potential mechanism in play. To quantify this effect, we measured the average distance worms moved when we flooded the microfluidics arena with 2.2 mM 1-oct, comparing the distances traveled in a 2-minute interval in control buffer and 1-oct. Worms significantly increased their locomotion in 1-oct, and returned to baseline rates 2-4 minutes after 1-oct withdrawal (Fig. 5E). Next, we created a simple behavioral simulation to determine whether this selective increase in locomotory activity could counteract the observed bias in reversal probabilities to achieve overall avoidance. The environment was modeled as a line divided into two equal compartments, representing the control buffer and the 2.2 mM 1-oct halves of the arena. Worms were modeled as one-dimensional particles, free to move along the full length of the line. They were allowed to reverse direction at random along the length of the line but were specified to reverse direction with fixed probabilities at the buffer/octanol interface, depending on which direction they approached from. Finally, the particles moved at different relative rates in the buffer and 1-oct compartments (Fig. 5F). Using our measured entry and exit reversal probabilities as inputs, this simulation showed that relative locomotory speed is a critical factor determining overall chemotaxis behavior, with a 60% speed increase sufficient to convert attraction to repulsion, given our observed interface reversal probabilities in the stripe chips (Fig. 5F). This simulation was highly simplified and did not account for the persistence of higher locomotion speeds once worms exited 1-oct (Fig. 5E), which could further potentiate repulsion since faster-moving exiting worms will be oriented away from the 1-oct zone, biasing their trajectories toward the further reaches of the buffer zone where they eventually slow. Importantly, these results extend the idea that 1-oct is encoded combinatorially, generating afferent inputs controlling at least three distinct aspects of locomotion: 1) promotion of reversals when encountering increased [1-oct], 2) promotion of reversals when encountering reduced [1-oct], and 3) increase of locomotory speed in the presence of increased [1-oct].

### Repulsive and attractive signaling competitively entrain the locomotory command neurons

To understand how the repulsive and attractive 1-oct signals affect locomotory reversals at a circuit level, we examined the activity pattern of the AVA reverse command interneuron in relation to 1-oct addition and withdrawal. First, it was necessary to develop a data analysis approach to detect and quantify sensory-driven patterns of AVA activation, given the well-documented patterns of spontaneous AVA activity state transitions (*23, 27, 54–59*) that could potentially obscure a sensory signal. To reduce the impact of these transitions, we averaged at least 6 recordings of AVA activity for each condition tested so that non sensory-associated AVA transitions would tend to average out with larger numbers of recordings, akin to spike-triggered averaging (*72–74*). We then calculated the ‘entrainment index’ (EI), as a measure of how strongly a response is positively or negatively correlated with 1-oct application, and quantified the probability that an observed correlation is greater than a random association (see Methods), setting a threshold of P<0.05 for non-random association. To validate this test, we analyzed ASH and AWC responses to 1-oct application, which were obviously highly correlated and anti-correlated, respectively. The ASH and AWC EIs showed the expected positive and negative values, with P <0.0015, indicating a less than one in six hundred chance that the correlations in neuronal responses with 1-oct applications were random (Fig. 6A, B). Applying this test to AVA activity, we observed strong inverse correlation with stimulant application at 0.22 mM 1-oct (the lowest tested concentration), and a significant negative EI value (Fig. 6C, D, H). This concentration was attractive and showed the lowest ASH:AWC activity ratio (Fig. 4C), suggesting that AWC may be driving AVA with minimal interference from ASH. At higher [1-oct] (0.66, 2.2 mM), AVA entrainment with 1-oct application was shifted positively (compared to 0.22 mM; Fig. 6C, D, H) and was not significantly different from random (P>0.05). At these concentrations, the ASH:AWC activity ratio is higher (Fig. 4C), consistent with the idea that a lack of entrainment may reflect interference between AWC and ASH signals impinging on AVA.

**Fig. 6.**
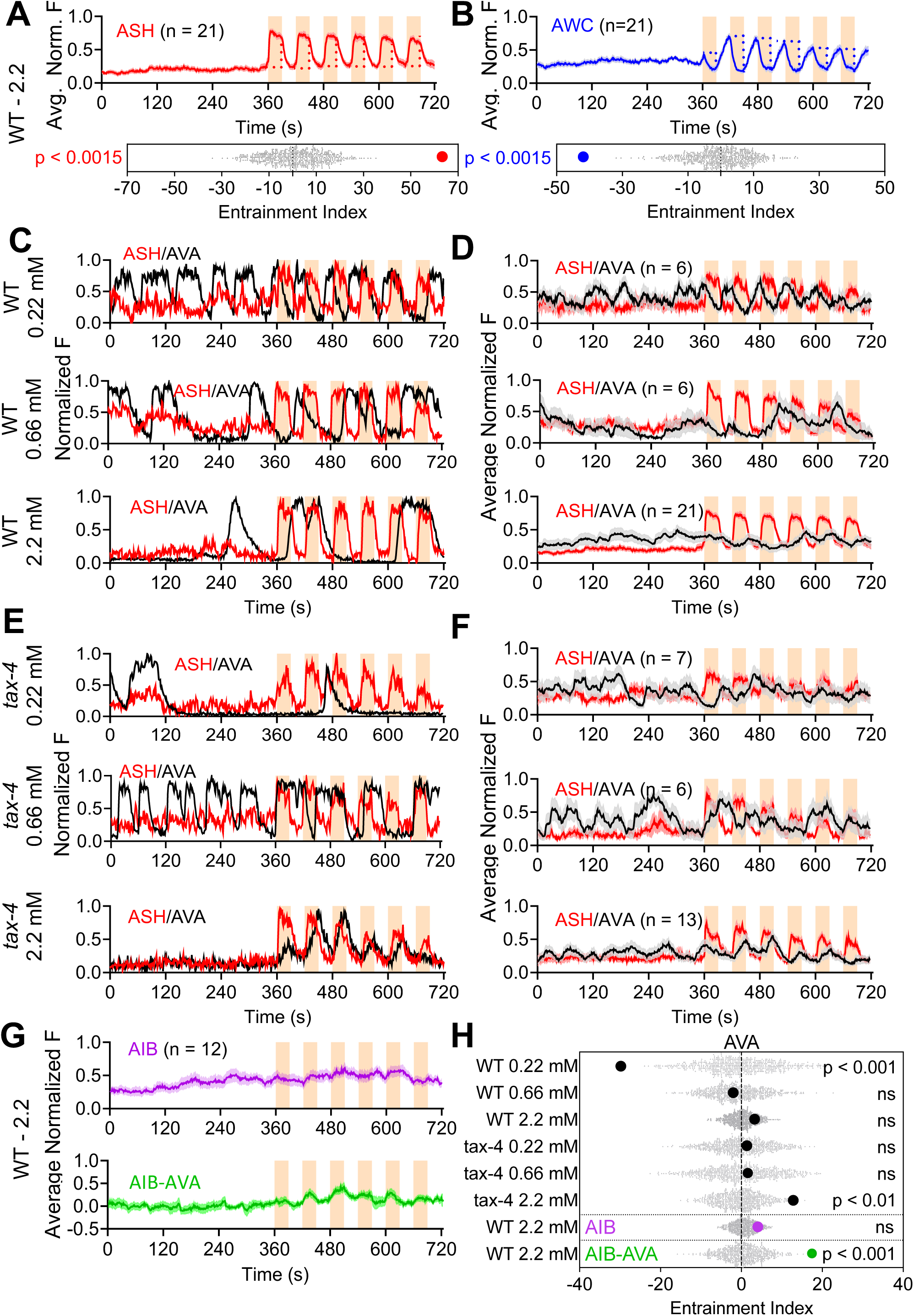
Entrainment of neuronal activity by 1-oct stimulation. (**A, B**) Validation of the entrainment index measurement using ASH (A) and AWC (B), showing very strong positive and very strong negative entrainment by 1-oct application, as expected. Gray dots indicate each randomized sum, while red and blue dots indicate the stimulus-specific sums of ASH and AWC, respectively (see Methods). (**C-F**) Superimposed ASH and AVA activity patterns after stimulation by 0.22, 0.66, and 2.2 mM 1-oct in wild type (C, D) and *tax-4* (E, F). Single worm traces (C, E) and averaged traces are shown (D, F, n ranges from 6-21 worms). (**G**) AIB traces (averaged); purple is unprocessed AIB signal, green is AIB minus AVA signal. (**H**) Entrainment indices for all recordings in C through G, with gray dots indicating randomized EIs and the bold dots represent the stimulus-specific EIs for AVA (black), unprocessed AIB (purple), and AVA-subtracted AIB (green). P values in A, B and H represent the probability that a randomized EI would be as great or greater than the stimulus-selected sum, with a significance cut-off of P<0.05.

To further support the idea that conflicting sensory signals may mask one another at the level of AVA, we measured EIs in TAX-4 animals. At the lowest [1-oct], the strong negative entrainment previously observed was abolished, confirming that negative entrainment is a function of TAX-4 signaling (Fig. 6E, F, H). At the intermediate [1-oct] (0.66 mM), *tax-4* loss of function mutation caused a slight positive shift in the EI, but did not lift it above the threshold for significant entrainment (P>0.05) (Fig. 6E, F, H). However, at the highest [1-oct], loss of TAX-4 positively shifted the EI into the significant range for positive entrainment (Fig. 6E, F, H). Interestingly, all of the calculated AVA EIs (Fig. 6H) correlate well with the overall chemotactic behavior in terms of sign (i.e negative entrainment predicts attraction, positive entrainment predicts repulsion), even where the strength of entrainment with the sensory input did not exceed our set threshold for non-random entrainment (see Discussion). Most importantly, these results imply that the reverse command interneurons under these assay conditions will not necessarily show overt entrainment to sensory stimuli even though they are receiving sensory inputs because sensory stimuli may be encoded as parallel antagonistic afferent inputs which cancel one another out, until the prevailing conditions permit one input to become substantially stronger than the other.

To investigate how the 1-oct sensory signals are transmitted to AVA, we examined the interneuron AIB, which is positioned to be an integrating hub in this system because it receives afferent inputs from ASH, AWC, and many other sensory neurons, and from RIM, which provides feedback about the motor state that is used in the regulation of sensory afferent pathways by corollary discharge (*54, 75, 76*). Initially, we did not observe significant entrainment of AIB activity with 2.2 mM 1-oct application (Fig. 6G). It is possible that immobilization and anesthetization may be affecting AIB responses to sensory activity and/or proprioceptive feedback from locomotion. However, it is also possible that motor feedback from RIM was obscuring the sensory signal. To address this possibility, we subtracted AVA activity, representing the motor state, from the AIB activity (AVA closely mirrors RIM), based on the observation that AIB activity can be modeled as the sum of convolutions of motor activity and sensory activity (*77*). This modified AIB signal showed significant entrainment with 2.2 mM 1-oct application which was positive in sign (Fig. 6 G and H), and greater than AVA’s entrainment at the same 1-oct concentration. These observations suggests that AIB serves as a sensory relay for the aversive inputs, but since aversive entrainment in AVA is weaker than that in AIB (presumably due to interference from attractive inputs), the site of integration between repulsive and attractive pathways must be downstream of AIB and either at or upstream of AVA.

### Combinatorial olfactory coding facilitates context-dependent behavioral flexibility

What benefit might *C. elegans* gain from encoding 1-oct combinatorially, simultaneously activating attractive and repulsive afferent input pathways? This strategy could create a simple and versatile mechanism to modulate the behavioral output to best match internal and external conditions, such as specific odorant concentration, the presence of other odorants, or interoceptive inputs such as hunger or stress. A combinatorial strategy allows response sensitivity and even valence to be altered by simply adjusting gains or synaptic outputs within each pathway (Fig. 7A, B), without requiring extensive functional reorganization of the downstream connections, or recruitment of complex cross-inhibitory circuitry. For example, in the laboratory, 1-oct is used as a repellant for *C. elegans*, but its presence in the environment may convey more complex information. 1-oct is toxic to worms (thus representing a risk), which presumably explains why *C. elegans* has evolved to avoid it, at least at higher concentrations. However, 1-oct is naturally found in decaying plant material (*78, 79*), so it may also signal the presence of a bacterial food source for worms (thus representing a reward as well). In the presence of non-toxic alternatives, other food sources are safer, but in the absence of alternatives, worms may risk toxicity to avoid starvation. Could combinatorial coding support the risk-reward decision making required of *C. elegans* when evaluating environmental 1-oct signals? Consistent with this idea, many attractive food signals, including isoamyl alcohol (IAA), suppress AWC activity (*35, 39, 61*), and could therefore decrease the activity of the attractive afferent pathway to disinhibit repulsion (Fig. 7A, B). We tested this hypothesis by conducting chemotaxis experiments at 0.66 mM 1-oct, normally an attractive concentration in our microfluidics assays, in the presence of isoamyl alcohol (IAA, 10^-4^ dilution, added to both the 1-oct and control buffer sides of the microfluidics arena). This concentration of IAA is strongly attractive in microfluidics assays (reference (*50*) and our unpublished data). Under these conditions, worms were strongly repelled by 1-oct (Fig. 7C). The probability of reversals when entering the 1-oct zone was increased (Fig. 7D), the probability of reversals when exiting the 1-oct zone was decreased (Fig. 7D), and the repulsion-inducing modulation of locomotory speed was also increased (Fig. 7E, F). Interestingly 0.66 mM 1-oct did not cause a speed increase on its own (unlike 2.2 mM, Fig. 5). Acute application of IAA also did not affect speed. However, 0.66 mM 1-oct in the presence of IAA caused a large speed increase (2-fold). Since IAA reduces AWC activity, this IAA effect is consistent with removal of AWC inhibition of 0.66 mM 1-oct-dependent speed modulation. These three quantitative changes in locomotory parameters will each shift the chemotaxis index in the negative direction, converting 1-oct attraction to repulsion.

**Fig. 7.**
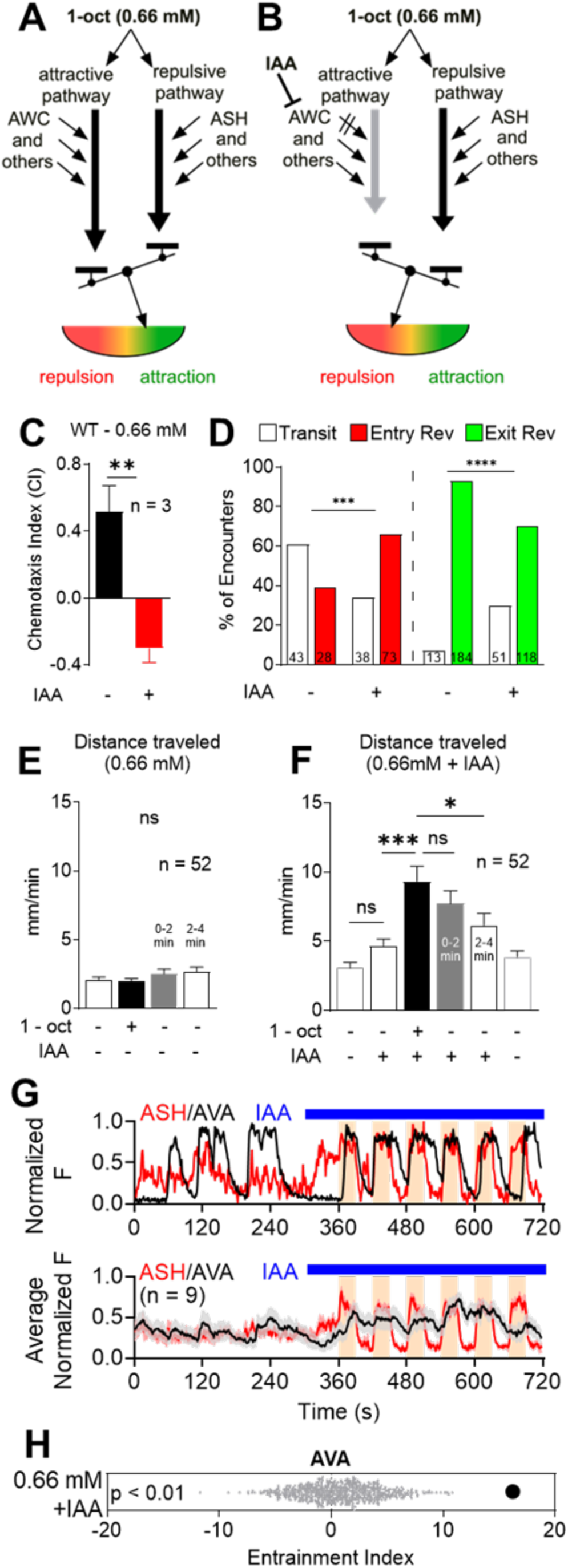
Balanced repulsion and attraction support context-dependent modulation of sensory responses. (**A, B**) Combinatorial coding-based mechanism for modulating sensory responses. (**A**) At lower 1-oct concentrations, the attractive afferent pathway is relatively strong, resulting in attraction. (**B**) Addition of IAA reduces AWC activity, weakening the attractive pathway relative to the repulsive pathway, and reversing the 1-oct valence. (**C**) Addition of 0.92 mM IAA reverses the valence of 1-oct (0.66 mM) from attractive to repulsive. (**D**) IAA biases reversal behavior (at the 1-oct/buffer interface) toward repulsion. Reversals upon entering 1-oct zone are increased (left side/red), and reversals upon exiting the 1-oct zone are decreased (right side/green). (**E-F**) Effect of IAA on locomotory activity. (**E**) 1-oct (0.66 mM) in the absence of IAA does not cause increased speed. (**F**) 1-oct (0.66 mM) in the presence of IAA causes markedly increased speed. (**G**) Single (upper) and averaged (lower) recordings of ASH and AVA before and after addition of IAA and 1-oct (0.66mM) as shown, demonstrating increased AVA entrainment to stimulus (compare to Fig-6D/middle). Blue bar indicates addition of IAA (in both control buffer and 1-oct solutions). (**H**) AVA entrainment index confirms that the presence of IAA causes AVA to become significantly entrained with the stimulus. ** in C indicates P<0.01, *t-test*; *** and **** in D indicate P<0.001, 0.0001 respectively, Fisher’s Exact Test; *, *** in F indicate P<0.05 and 0.001, respectively, ANOVA and Tukey’s multiple comparison test.

In network recordings, the presence of IAA greatly increased the sensory entrainment of AVA, positively correlating with stimulus application. This effect suggests that IAA will potentiate 1-oct avoidance by klinokinesis, or in other words, increasing the probability that worms will execute a reversal when they experience positive d[1-oct]/dt. These results provide proof-of-principle that rebalancing the afferent input pathways to reduce attraction and promote repulsion can control chemotaxis in an ethologically relevant manner and demonstrate the potential utility and flexibility of the combinatorial coding strategy.

## Discussion

We have investigated the response of *C. elegans* to the odorant 1-oct, using a combination of microfluidics-based behavioral analysis and whole-network Ca^++^ imaging. Ca^++^ imaging using the NeuroPAL system (*30*) showed that 1-oct activates at least 11 different sensory neurons including neurons associated with chemorepulsion and chemoattraction. 1-oct activates the ASH and AWC sensory neurons, with ON- and OFF-response kinetics, respectively. Paradoxically, this pattern suggests that sensory signals are being generated to simultaneously promote and inhibit locomotory reversals upon both 1-oct onset and offset. Based on this observation, we hypothesized that 1-oct may simultaneously activate repulsive and attractive afferent pathways, with the ultimate chemotactic outcome being determined by the balance between them. We tested this hypothesis using several approaches to selectively modulate the input pathways, such as varying 1-oct concentration, mutating key proteins in sensory signaling pathways, and selectively inactivating ASH and AWC chemogenetically. We observed that decreasing the repulsive drive unmasked attractive responses, and vice versa, indicating that balancing attractive and repulsive sensory afferent pathways control wild type 1-oct responses over at least a ten-fold concentration range. This novel insight highlights the value of the whole-brain approach (enabled by the NeuroPAL system) for studying the network dynamics underlying sensory driven behaviors.

Three distinct locomotory transitions appear to be important for the overall chemotactic outcome: reversals stimulated upon entering the 1-oct zone (‘entry reversals’), reversals stimulated upon exiting the 1-oct zone (‘exit reversals’), and increased locomotory activity within the 1-oct zone (‘speed increase’) (Fig. 8A). Based on numerical simulations, speed increase was the most important factor: reversal patterns at the highest 1-oct concentration clearly favored attraction but increased locomotory activity within the 1-oct zone overcame this bias and caused repulsion. Our results suggest that both repulsive and attractive afferent pathways may be involved in regulating these key locomotory transitions (Fig. 8A). Entry reversals were reduced in *osm-9* mutants but increased in *tax-4* mutants. Exit reversals were reduced in *tax-4* mutants (but not affected in *osm-9*). Speed increase was dependent on 1-oct concentration (akin to the activity of the repulsive input pathway) and potentiated under conditions in which depress the activity of AWC (a key sensory neuron in the attractive pathway).

**Fig 8.**
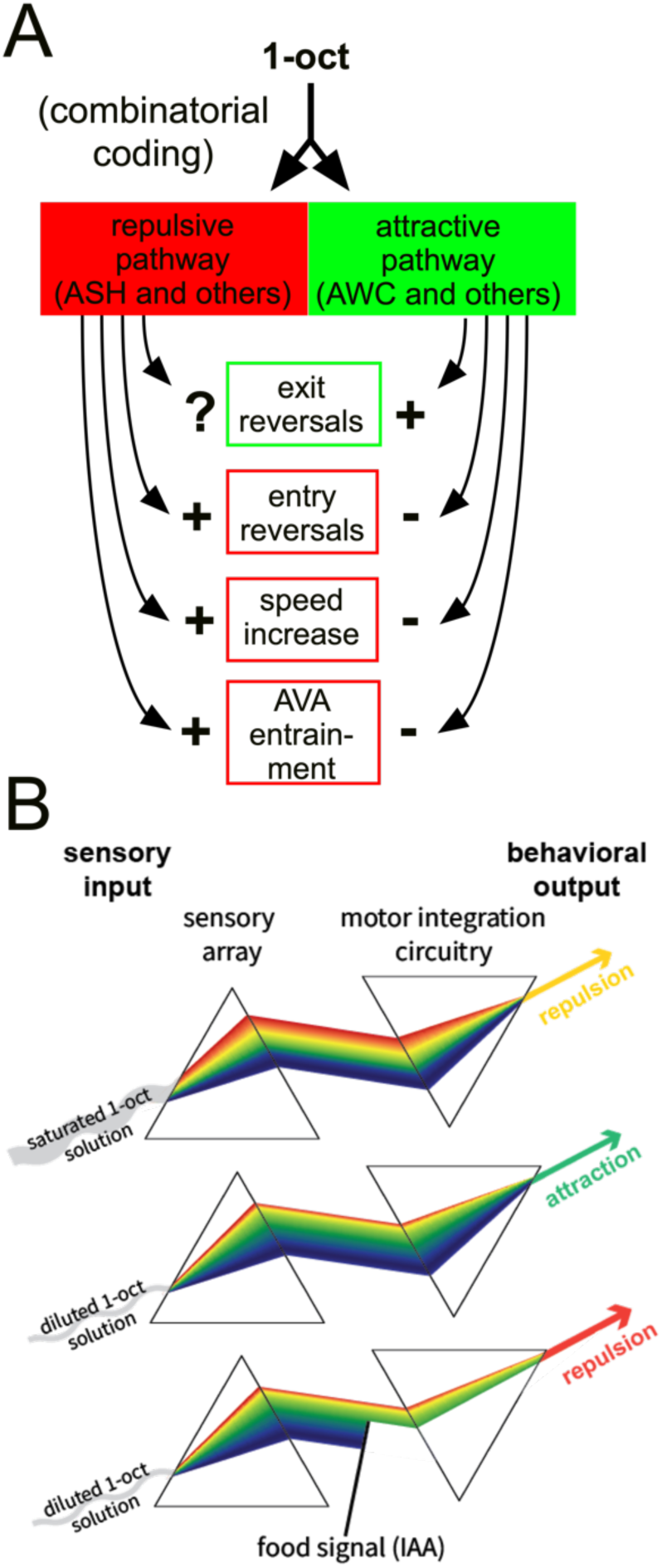
Combinatorial coding of 1-oct as an attractant and a repellant, simultaneously, facilitates sensory motor coupling and context-dependent modulation in *C. elegans*. **(A)** 1-oct simultaneously activates sensory neurons in the periphery that mediate both repulsion (ASH and others) and attraction (AWC and others), such that repulsive and attractive afferent pathways become active concurrently. These inputs are then integrated centrally to control at least three key locomotory parameters (exit reversals, entry reversals, and locomotory speed), and the activity state of the reverse command neurons. **(B)** An analogy based on refraction on light through two prisms highlights the potential versatility of olfactory combinatorial coding in *C. elegans*. An individual odorant activates many chemosensory neurons to produce multiple distinct afferent inputs propagating through the downstream circuitry simultaneously, analogous to the first prism separating white light into its component colors (top panel, left). The motor integration circuitry formulates a locomotory response by integrating the active afferent pathways, analogous to the second prism recombining the incoming light (top panel, right). Because chemosensory neurons have different odorant affinities, the specific outputs of the sensory array and motor integration circuit may be concentration dependent. In this study, ASH (coded in red) is a low affinity sensor, while AWC (coded in blue) is a high affinity sensor. Reducing the 1-oct concentration biases the output of the sensory array toward AWC (middle panel, left), and the motor integration circuitry uses this information to produce an attractive locomotory response (middle panel, right). This coding strategy produces a simple framework for modulation. The behavioral outcome may be influenced by simply shifting the balance of the afferent pathways, much like an optical filter placed in a light path can selectively eliminate specific wavelengths (lower panel) to change the color of the transmitted light. In this study, IAA was added to the diluted 1-oct solution, inhibiting AWC, such that the afferent pathways were re-balanced toward the ASH end of the spectrum, and repulsion was restored. This ‘filtering’ effect could, in principle, take place peripherally or centrally, and could subserve modulation due to the presence of other odorant molecules, monoamine/neuropeptide modulators serving as interoceptive signals, and effectors of experience-dependent plasticity.

Our chemogenetic data provide strong evidence that ASH and AWC contribute significantly to the repulsive and attractive input pathways, respectively. However, we cannot rule out the possible contributions of other sensory neurons. The promoters used for transgenic expression of HisCl1 are active in a few other neurons (i.e. P*sra-6* is active in ASI and *ceh-36* is active in ASE). However, these neurons showed little to no 1-oct responsiveness. For the repulsive pathway, the minimal effects due to loss of OSM-9 suggest that repulsive drive may also be provided by TAX-4-positive neurons (with the caveat that some ASH signaling remains in *osm-9* mutants) (*38, 67*). AWB and BAG are candidates based on their consistent 1-oct responses and the observation that AWB contributes to avoidance responses to volatile 1-oct (*34*). For the attractive pathway, chemogenetic inactivation of AWC produced a modest effect relative to loss of TAX-4, implying there may be other TAX-4-dependent neurons contributing. We also cannot rule out a role for AWA, an OSM-9-dependent neuron associated with chemoattraction to soluble compounds. More generally, any sensory neuron which consistently responds to 1-oct addition or withdrawal has the potential to contribute to the behavioral response. Systematic mapping of sensory neuron contributions, individually and in combination, to the locomotory transitions and sensory-driven activity patterns in the downstream neural circuitry will be important to fully understand how *C. elegans* generates its behavioral response to 1-oct, and eventually other odorant classes.

Our results show how combinatorial and labeled line coding schemes may functionally interact. 1-oct is encoded combinatorially, in that it activates multiple sensory neurons (our results, and (*39*)), principally ASH and AWC. However, these neurons have long been regarded to represent repulsive and attractive labeled lines, respectively (*33, 35, 38, 80*). Thus, the initial combinatorial representation of 1-oct superimposes on the underlying labeled line architecture to dictate the final chemotactic outcome. The novel finding here is that multiple labeled lines become activated simultaneously in the context of a combinatorially-coded odorant. Previous studies have shown that antagonistic sensory inputs can contribute to chemosensory responses in *C. elegans* (*81–84*), but this study is the first to show in detail how conflicting inputs interact at the behavioral and circuit levels. The labeled lines independently control multiple locomotory parameters, in opposite directions, and the conflict is resolved by the locomotory system which is sensitive to the balance of the two pathways.

The compact size of the *C. elegans* nervous system makes it feasible to characterize at a cellular and molecular level where and how these afferent pathways converge and interact. Both pathways ultimately impinge on the AVA locomotory command interneurons, which drive locomotory reversals (*85, 86*). Because AVA sensory entrainment varies smoothly as a function of the relative activity levels of the two pathways, they probably do not strongly cross-inhibit one another, which would likely produce much steeper transitions (*87*). Which neurons are likely to integrate the attractive and repulsive inputs? Based on the connectome and previous functional studies, the AIB interneuron is a logical candidate. AIB is a first layer interneuron that receives extensive inputs from sensory neurons including ASH and AWC, strong synaptic inputs from other first layer interneurons (AIZ, AIA), and electrical and chemical synapses from locomotory command neurons AVE and RIM. AIB sends projections to the locomotory command neurons including AVA, RIM, and AVB (*63, 88*). Functionally, AIB serves as an integrating hub for multiple sensory inputs including ASH and AWC (*89, 90*). However, AIB does not appear to play the role of an integrating hub upstream of AVA for 1-oct. Prior to sensory stimulation, AIB activity closely followed reverse command interneuron activity, as has been observed previously (*23, 54*). Upon stimulation, AIB still followed the reverse command interneurons, but subtracting the reverse command signal from the AIB excitation pattern revealed that AIB was significantly entrained to the repulsive sensory input. AVA did not show this repulsive entrainment under the same conditions, therefore AIB cannot be the sole locus for integration. One intriguing possibility is that integration takes place in AVA itself, since it receives strong synaptic inputs directly from ASH, and from SAA, which mirrors the activity pattern of AWC during stimulation.

This coding scheme facilitates olfactory plasticity because the balance of the pathways, evaluated centrally, can determine odor valence. This plasticity differs from what has previously been described, where odorant responses are modulated at the periphery through lateral interactions among sensory afferents (*20, 35*) or reprogramming of individual sensory neuron responses (*47*). The broad versatility of this coding scheme can be appreciated by considering combinatorial coding and integration of inputs as analogous to the refraction of light through two prisms (Fig. 8B). First, the chemosensory neuron array in *C. elegans* creates a multi-factorial representation of the sensory stimulus, (i.e. the first prism), differentiating inputs from multiple sensory neurons, each with different downstream behavioral correlates. Downstream, the different components of this representation will eventually converge and become integrated into a coherent behavioral response by the motor command circuitry (i.e. the second prism). Because the sensory neurons have different concentration-response profiles, worms can easily generate differential behaviors dependent on stimulus concentration: as concentration is reduced, the input from the sensory layer will be skewed toward the higher-affinity components (Fig. 8B). In this study, reduced [1-oct] favored the AWC-driven response (higher affinity) over the ASH-driven response (lower affinity), converting the 1-oct response from repulsion to attraction. This architecture also provides straightforward avenues for modulation. Multiple sensory inputs may be integrated if the additional odorants modulate the initial sensory representation, either additively (via ON neurons) or subtractively (via OFF neurons Fig. 8B). Here, the addition of IAA restored repulsion at lower concentrations by reducing the contribution of AWC. In the two-prism analogy, IAA is equivalent to a selective optical filter placed in the light path, skewing the overall balance back toward ASH. Other forms of plasticity, such as monoamine and neuropeptide modulation or experience dependent plasticity, may also be easily incorporated into this scheme at the sensory- or interneuron level. In a complex natural environment, where multiple odorants, interoceptive inputs, and experience-dependent plasticity are all acting simultaneously on a limited set of neurons, potential interference among modulatory signals might be avoided by funneling all the modulatory influences through one major repulsive pathway and one major attractive pathway.

## Materials and Methods

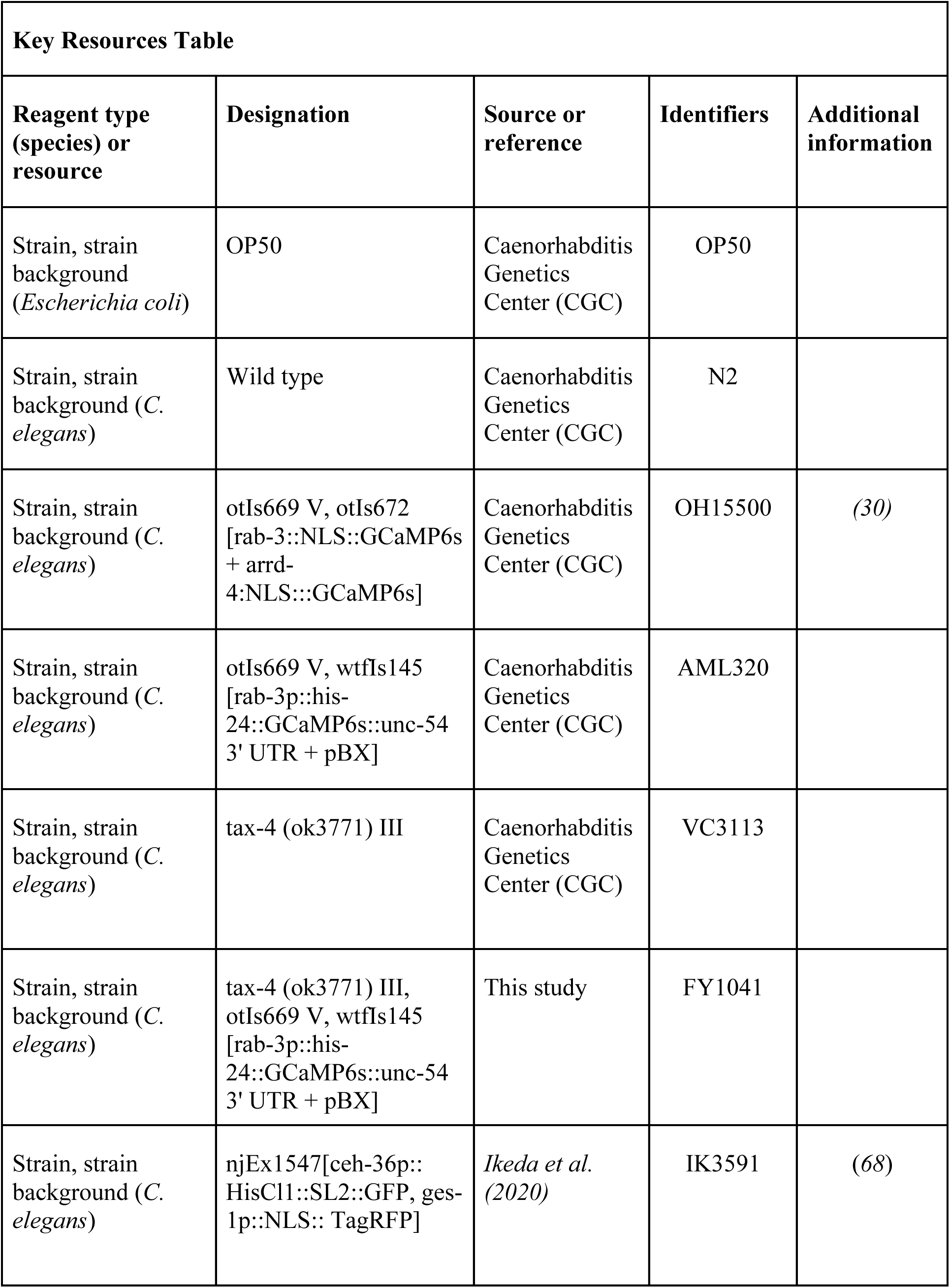
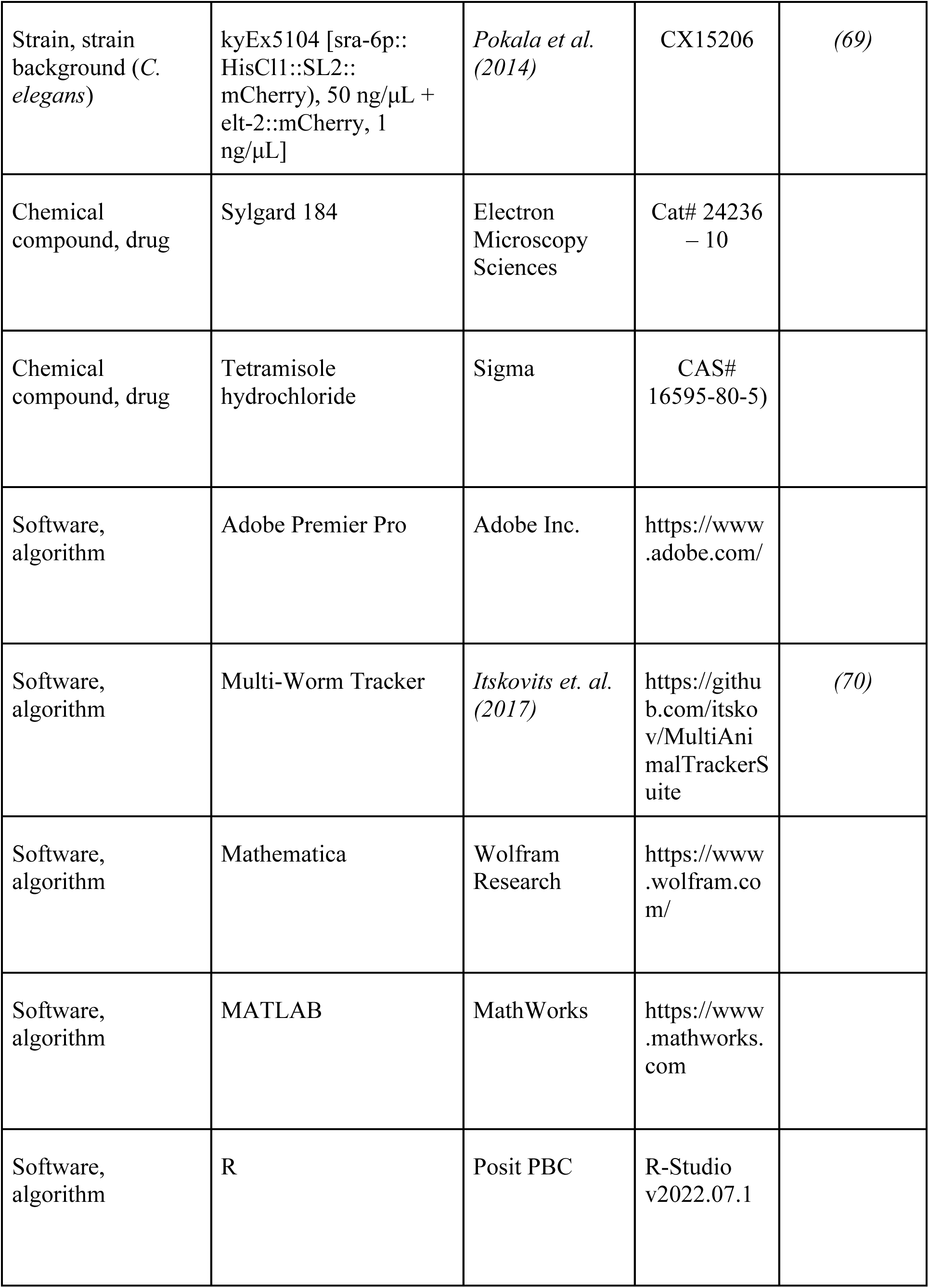
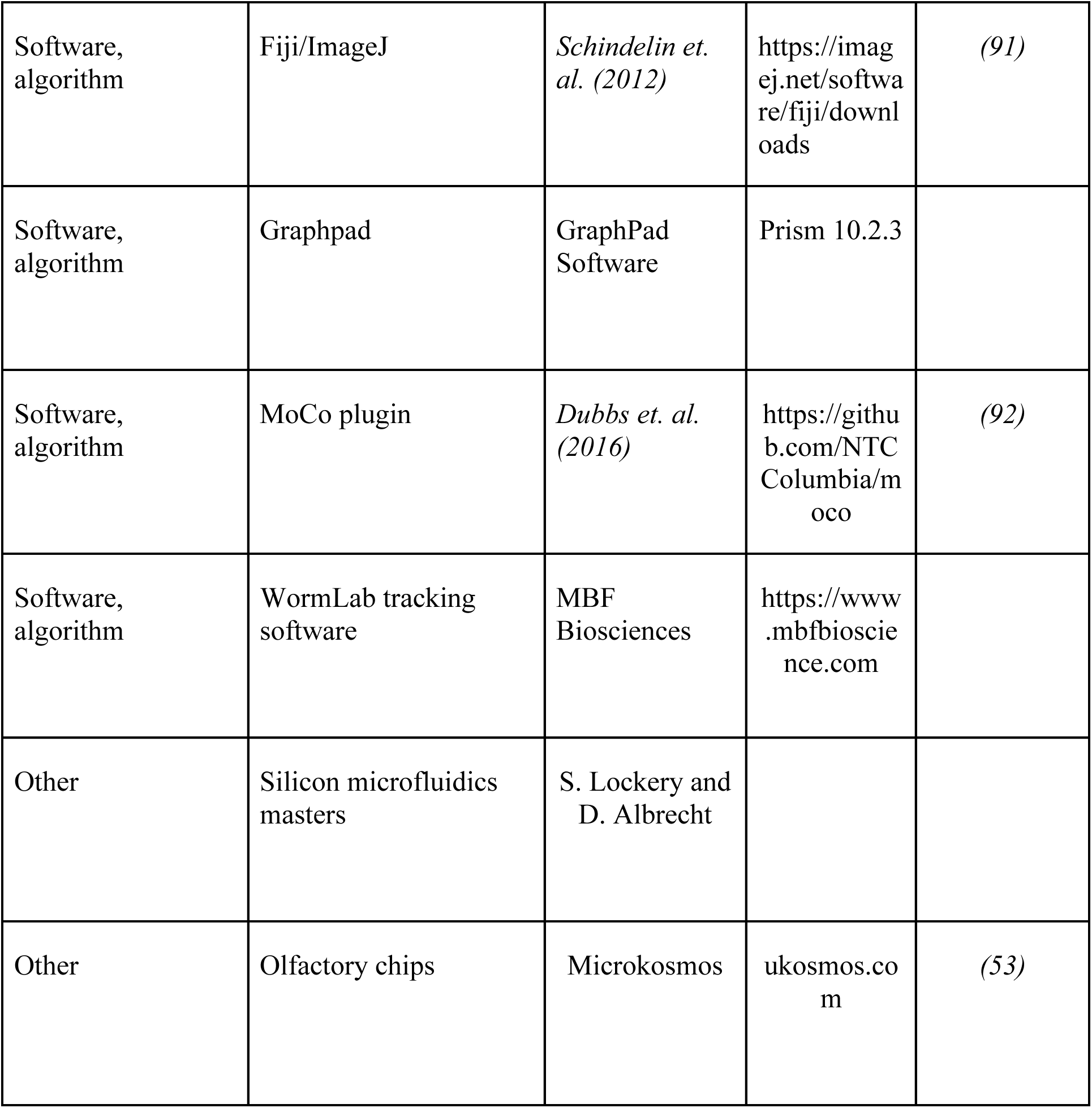

| Key Resources Table |  |  |  |  |
| --- | --- | --- | --- | --- |
| Reagent type (species) or resource | Designation | Source or reference | Identifiers | Additional information |
| Strain, strain background (*Escherichia coli*) | OP50 | Caenorhabditis Genetics Center (CGC) | OP50 |  |
| Strain, strain background (*C. elegans*) | Wild type | Caenorhabditis Genetics Center (CGC) | N2 |  |
| Strain, strain background (*C. elegans*) | otIs669 V, otIs672 [rab-3::NLS::GCaMP6s + arrd-4::NLS::GCaMP6s] | Caenorhabditis Genetics Center (CGC) | OH15500 | (30) |
| Strain, strain background (*C. elegans*) | otIs669 V, wtfIs145 [rab-3p::his-24::GCaMP6s::unc-54 3' UTR + pBX] | Caenorhabditis Genetics Center (CGC) | AML320 |  |
| Strain, strain background (*C. elegans*) | tax-4 (ok3771) III | Caenorhabditis Genetics Center (CGC) | VC3113 |  |
| Strain, strain background (*C. elegans*) | tax-4 (ok3771) III, otIs669 V, wtfIs145 [rab-3p::his-24::GCaMP6s::unc-54 3' UTR + pBX] | This study | FY1041 |  |
| Strain, strain background (*C. elegans*) | njEx1547[ceh-36p::HisC11::SL2::GFP, ges-1p::NLS:: TagRFP] | *Ikeda et al. (2020)* | IK3591 | (68) |
| Strain, strain background (*C. elegans*) | kyEx5104 [sra-6p::HisC11::SL2::mCherry), 50 ng/μL + elt-2::mCherry, 1 ng/μL] | *Pokala et al. (2014)* | CX15206 | (69) |
| Chemical compound, drug | Sylgard 184 | Electron Microscopy Sciences | Cat# 24236 – 10 |  |
| Chemical compound, drug | Tetramisole hydrochloride | Sigma | CAS# 16595-80-5) |  |
| Software, algorithm | Adobe Premier Pro | Adobe Inc. | https://www.adobe.com/ |  |
| Software, algorithm | Multi-Worm Tracker | *Itskovits et. al. (2017)* | https://github.com/itskov/MultiAnimalTrackerSuite | (70) |
| Software, algorithm | Mathematica | Wolfram Research | https://www.wolfram.com/ |  |
| Software, algorithm | MATLAB | MathWorks | https://www.mathworks.com |  |
| Software, algorithm | R | Posit PBC | R-Studio v2022.07.1 |  |
| Software, algorithm | Fiji/ImageJ | *Schindelin et. al. (2012)* | https://imagej.net/software/fiji/downloads | (91) |
| Software, algorithm | Graphpad | GraphPad Software | Prism 10.2.3 |  |
| Software, algorithm | MoCo plugin | *Dubbs et. al. (2016)* | https://github.com/NTC-Columbia/moco | (92) |
| Software, algorithm | WormLab tracking software | MBF Biosciences | https://www.mbfbioscience.com |  |
| Other | Silicon microfluidics masters | S. Lockery and D. Albrecht |  |  |
| Other | Olfactory chips | Microkosmos | ukosmos.com | (53) |

### Worm maintenance and strains

*C. elegans* were maintained at 20°C on NGM agar plates and fed *E. coli* OP50 bacteria. All experiments were conducted at 20-22°C (*93, 94*). The following strains were utilized:

**N2**: *Wild-type*.

**OH15500**: otIs669 V, otIs672 [rab-3::NLS::GCaMP6s + arrd-4:NLS:::GCaMP6s] (*30*).

**AML320**: otIs669 V, wtfIs145 [rab-3p::his-24::GCaMP6s::unc-54 3’ UTR + pBX].

**VC3113**: tax-4 (ok3771) III.

**CX10**: osm-9 (ky10) IV.

**FY1041**: tax-4 (ok3771) III, otIs669 V, wtfIs145 [rab-3p::his-24::GCaMP6s::unc-54 3’ UTR + pBX].

**IK3591**: njEx1547[ceh-36p::HisCl1::SL2::GFP, ges-1p::NLS::TagRFP]

**CX15206**: kyEx5104[sra-6p::HisCl1::SL2::mCherry, 50 ng/μL + elt-2::mCherry, 1 ng/μL]

Strains were obtained from the Caenorhabditis Genetics Center (CGC), funded by NIH (P40 OD010440). VC3113 was crossed into AML320 to generate FY1041, with the presence of *tax-4* verified by PCR. Strains were allowed for three generations of growth before experiments.

### Microfluidics device fabrication and behavioral assays

All behavioral studies were conducted in microfluidic devices following standard protocols (*49–51*). Devices were fabricated by thoroughly mixing polydimethylsiloxane (PDMS) and a curing agent in a 10:1 ratio by weight (Sylgard 184, Electron Microscopy Sciences Cat. 24236 – 10), degassing the mixture under vacuum, and then pouring it onto a silicon mold at ̴̴ 5 mm height. PDMS was cured by incubating at 65°C for at least 3 hours. Olfactory chips were obtained from Microkosmos (ukosmos.com).

### Chemotaxis

Chemotaxis assays utilized a ‘two-chamber’ device dividing the arena into equal chambers for stimulus and control buffer (S-basal [100mM NaCl, 5mM g K_2_HPO_4_, 45mM KH_2_PO_4_, pH = 6]) (*49*). Worm behavior was recorded using AmScope microscope cameras (MD 500, MU 500-HS, AmScope, Irvine, CA), and videos were processed using Adobe Premier Pro 2024 (Adobe Inc., San Jose, CA). 1-2 day old adults were picked and washed with S-basal and loaded into the microfluidics arena. Animals were acclimatized in the arena for 5 minutes; assays were performed between 20-40 minutes from starvation onset while they were in area-restricted search foraging mode (*52*). In the original microfluidics protocols (*50, 51*) 0.01% w/v xylene cyanol (0.1mg/ml) dye was used to verify clean separation between different domains within the microfluidics device. However, worms show clear attraction to 0.01% xylene cyanol (data not shown), indicating that the presence of dye introduces an additional olfactory stimulus. Therefore, we modified the protocol to test the flow pattern using dye before and after the assay but remove the dye during the assay (Fig 1A). In control trials using dye, we verified that flow patterns stay stable for up to 90 minutes once properly established. If the flow pattern in the post-data collection dye test was compromised, data were rejected. Histamine incubation was performed as previously described (*69*). Chemotaxis index was calculated between 10-20 minutes of stimulus onset, using the equation: Chemotaxis Index (CI) = (Number of worms in stimulus – Number of worms in buffer)/Total number of worms, positive and negative indices representing attraction and aversion respectively.

### Distance traveled

Locomotory distance-traveled assays used the ‘two-chamber’ device (*49*) with both chambers containing the same solution. Worms were exposed sequentially to buffer (5 mins), stimulus (5 mins), and buffer (10 mins). A complete exchange of solutions takes approximately 1 minute. Data were collected 90 seconds after solution change to ensure complete exchange. Distance traveled per minute was recorded over 2 minutes in buffer (pre-stimulus), in 1-oct solution (stimulus), and in successive 2-minute intervals after 1-oct withdrawal (post-stimulus). Worm tracks were manually traced and measured using Adobe Premiere Pro 2024 and Graphic for Mac (Autodesk, San Francisco, CA).

Worm locomotion simulation was written in Mathematica using AI-enabled chat (Mathematica, Princeton NJ). In the simulation, 20 one-dimensional dots, representing worms, were allowed to travel a one-dimensional line, representing behavior arena. The line was divided equally into two zones, representing buffer and stimulus, and dots moved freely within the line with random reversal frequency. Dot speed increased in the stimulus zone, as indicated, and reversal frequency at the interface was specified as observed from behavioral assays. 10 simulations were performed and averaged for each speed differential shown in Fig. 5F.

### Reversals

Analyses of locomotory reversals were performed using larger ‘stripe’ chips (*50*), which provided a greater working area and facilitated automated video tracking by reducing the frequency of collisions between worms. These chips were usually configured with 3 zones of equal size: a stripe of 1-oct down the middle flanked on either side by buffer, although for some experiments, the 1-oct was placed on one side. Video recordings of worm behavior were analyzed using the Multi-Worm Tracker (*95*) using MatLab (Mathworks, Natick, MA), or the WormLab tracker (MBS Biosciences, Williston VT), with manual analysis of individual tracks to quantify reversals.

### Whole-brain calcium imaging

Neuronal activity was examined by measuring calcium transients detected by fluorescence change (GCaMP6s, OH15500 (*30*), AML320 (*96*)). Calcium imaging was conducted as previously described (*23, 53, 54*). Animals were anesthetized with 1 mM tetramisole hydrochloride (Sigma, CAS 16595-80-5) in S-basal buffer and imaged in PDMS olfactory microfluidic devices, which trap the worm in a chemical delivery system and provides inlets to deliver either buffer or stimulus to the nose using laminar flow (*53*). Head-ganglia neurons were recorded using Leica confocal microscope (SP8) at 0.6 - 0.8 Hz rate. Odorants were diluted in S-basal. Each worm was recorded for 12-14 minutes within 20-40 minutes from starvation onset, starting with 6 minutes of buffer exposure followed by 6-7 30-second pulses of stimulus with 30-second intervals. Calcium imaging was followed by immediate recording of NeuroPAL color markers. For +IAA experiments, IAA delivery to the nose was started after 5 minutes of recording (1 minute before the first stimulus application) and IAA was was present in the stimulus and buffer channels thereafter. For concentration-response of ASH and AWC experiments, three 0.22 mM, two 0.66 mM and two 2.2 mM 1-oct pulses were given serially to the same worm; repeated application produced identical response amplitudes.

Recorded image files were stacked and analyzed using Fiji/ImageJ (*91*) followed by motion correction with ‘MoCo’ plug-in (*92*). Neuron identity was manually assigned and matched with calcium response using guidelines described by Yemini and Hobert (*30*). Neurons with ambiguous identities were excluded from the analysis (Fig. 2). Individual neuron activity was then normalized and re-sampled, before averaging, by ‘Min-Max normalization’ and ‘Linear interpolation’ respectively, using custom R-scripts (*97*).

### Data analyses and statistics

Statistical analyses were performed using Prism v10.2.1 (GraphPad Software Inc., La Jolla, CA), R v4.3, R-Studio v2022.07.1 (Posit PBC, Boston, MA), MatLab vR2023A (MathWorks, CA) and Microsoft Excel 2016. Chemotaxis assays were repeated 3-5 times and analyzed by *Student’s t-test* (un-paired). Pooled data from distance-traveled assays were analyzed by one-way ANOVA with post-tests. Reversal assays were repeated 3 times, data were pooled, and analyzed using Fisher exact tests.

Entrainment indices (EIs) were calculated using averaged, normalized calcium responses. First, area-under-curve (AUC) values were calculated for all possible 30s epochs by summing the Ca^++^ signal intensities over 30s after first subtracting the corresponding initial Ca^++^ signal intensity. EIs were calculated by adding six AUCs; stimulus-dependent EIs were calculated by adding the six AUCs corresponding to stimulus application intervals, while random EIs were calculated by adding six randomly-chosen AUCs. The significance of stimulus entrainment was estimated as the probability that the stimulus-dependent EI would exceed any given random EI, with probabilities < 0.05 considered significant (i.e. P<0.05). A P value <0.0015 indicated that the experimental EI was larger (or smaller) than all possible random EIs within the dataset. EIs for AVA were calculated using a 10-second lag, across all conditions, because, where obvious sensory entrainment was seen, AVA signals showed consistently slowed responses compared to sensory signals. No lag was incorporated for sensory and first-layer interneurons (ASH, AWC, AIB).

Neuronal activity correlations were calculated using Spearman’s rank correlation test and corrected with Fisher’s z-transformation before testing statistical significance (*98–100*), using custom R scripts.

## Acknowledgments

We would like to thank Shawn Lockery and Dirk Albrecht for invaluable help and materials for the microfluidics behavioral assays, the *C. elegans* Genetics Center for strains (funded by NIH P40 OD010440), Ikue Mori for IK3591, Cori Bargmann for CX15206, Tomer Avidor-Reiss and Qian Chen for assistance with confocal microscopy,

## Author contributions

Conceptualization: MZHS, BAB, WGR, RM. Software: BAB. Methodology: MZHS, BAB, CAE, LGJ. Investigation: MZHS, WGR, CAE, BNS, LA, JK, AM, ARM, RM, BAB. Data visualization: MZHS, WGR, BAB. Supervision: BAB, RM. Writing (original draft): MZHS, BAB. Funding acquisition BAB, RM, WGR.

## Data and materials availability

Dataset and code are available for download at Zenodo.org (https://doi.org/10.5281/zenodo.13328278).

